# A novel and specific regulator of neuronal V-ATPase in *Drosophila*

**DOI:** 10.1101/2021.02.08.430253

**Authors:** Amina Dulac, Abdul-Raouf Issa, Jun Sun, Giorgio Matassi, Baya Chérif-Zahar, Daniel Cattaert, Serge Birman

**Affiliations:** Genes Circuits Rhythms and Neuropathology, Brain Plasticity Unit, CNRS, ESPCI Paris, PSL Research University, 10 rue Vauquelin, F-75005 Paris, France; Dipartimento di Scienze Agroalimentari, Ambientali e Animali, University of Udine, Udine, Italy and Université de Picardie Jules Verne, CNRS-UMR 7058 “Ecologie et Dynamique des Systèmes Anthropisés” (EDYSAN), Amiens, France; INCIA - Institut de Neurosciences Cognitives et Intégratives d’Aquitaine, CNRS, Université de Bordeaux, PAC Talence, allée Geoffroy Saint-Hilaire, F-33615 Pessac cedex, France

**Author notes:** Sussex Neuroscience, School of Life Sciences, University of Sussex, Biology Road, Brighton BN1 9QG, UK. Neurodevelopment Laboratory, School of Biosciences, University of Birmingham, Birmingham B15 2TT, UK.

**Keywords:** CG31030/VhaAC45L, neuronal V-ATPase, synaptic vesicle acidification, quantal size, *Drosophila melanogaster*

## Abstract

The V-ATPase is a highly conserved enzymatic complex that ensures appropriate levels of organelle acidification in virtually all eukaryotic cells. While the general mechanisms of this proton pump have been well studied, little is known about the specific regulations of neuronal V-ATPase. Here, we studied CG31030, a previously uncharacterized *Drosophila* protein predicted from its sequence homology to be part of the V-ATPase family. We found that this protein is essential and apparently specifically expressed in neurons, where it is addressed to synaptic terminals. We observed that CG31030 co-immunoprecipitated with V-ATPase subunits, in particular with ATP6AP2, and that synaptic vesicles of larval motoneurons were not properly acidified in *CG31030* knockdown context. This defect was associated with a decrease in quantal size at the neuromuscular junction, severe locomotor impairments and shortened lifespan. Overall, our data provide evidence that CG31030 is a specific regulator of neuronal V-ATPase that is required for synaptic vesicle acidification and neurotransmitter release.

## Introduction

Many cellular processes require a specific electrochemical environment for proper functioning, such as post-translational modifications of proteins in the Golgi apparatus, lysosomal degradation, endosomal ligand-receptor dissociation, or hormone concentration (reviewed in Forgac, 2007). Eukaryotic cells use a highly conserved proton pump, called the vacuolar H^+^-ATPase (V-ATPase), to achieve the adequate level of acidity in different cellular compartments (Saroussi and Nelson, 2009). This large enzymatic complex must be tightly regulated, as it is essential for it to be localized on the right membrane, and to fit the different pH ranges specific to each organelle and cell type.

In neurons, the V-ATPase plays a crucial role at the synapse, being responsible for acidifying synaptic vesicles and thus providing the driving force for neurotransmitter loading (Moriyama et al., 1992). Recently, neuronal V-ATPase has also gained interest in the context of aging and neurodegenerative diseases, as its dysregulation, and resulting impairment of the autophagy-lysosomal pathway, have been linked to several pathologies such as Alzheimer and Parkinson disease (Colacurcio and Nixon, 2016; Collins and Forgac, 2020). If the core mechanism of the proton pump is now well understood, the regulations conferring the cell-specific functions of neuronal V-ATPase remain largely unknown, considerably limiting its potential use as a therapeutic target.

The V-ATPase complex is composed of a cytoplasmic domain (V_1_) and a membrane-bound domain (V_0_). The V_1_ domain contains the catalytic unit responsible for ATP hydrolysis. The energy resulting from this reaction powers a rotational molecular motor spanning from V_1_ to V_0_, allowing protons to cross membranes through the port contained in V_0_ (Vasanthakumar and Rubinstein, 2020). The assembly of V_1_ to V_0_ is necessary for the pump to function, and reversible dissociation of the two domains has been shown to occur as a way to regulate V-ATPase activity (Collins and Forgac, 2020). Though the core mechanism stays the same, one V-ATPase can differ from another by its composition. In vertebrates, as well as in *Drosophila*, V_0_ is made of five subunits (*a*, *c*, *c’’*, *d* and *e*), while V_1_ contains eight subunits (A, B, C, D, E, F, G and H) (Allan et al., 2005). Each subunit can have several paralogs encoded by different genes, and each gene can produce several isoforms, allowing many different possible combinations to form the full V-ATPase complex. These differences of composition can also have regulatory effects on the complex, both on its localization and on its functional properties (Vasanthakumar and Rubinstein, 2020).

The V-ATPase can also be regulated by two accessory subunits, ATP6AP1/Ac45 and ATP6AP2/PRR (Jansen and Martens, 2012). Both proteins are found in the nervous tissue but are also required in other organs, and their mutation has been linked to cognitive impairments as well as systemic symptoms like immunodeficiency or hepatopathy (Jansen et al., 2016; Cannata et al., 2018). These two subunits interact directly with the V_0_ domain and are believed to promote assembly of the membrane and soluble regions of the V-ATPase complex (Abbas et al., 2020). Another potential accessory subunit exists in vertebrates, named ATP6AP1-like (ATP6AP1L), which is homologous to ATP6AP1 and has not yet been functionally characterized. *Drosophila* possesses identified homologs of ATP6AP1/Ac45 and ATP6AP2/PRR, named VhaAC45 and ATP6AP2, respectively. These proteins also seem to contribute to assembly of the V-ATPase in fly tissues (Schoonderwoert and Martens, 2002a; Guida et al., 2018).

In this study, we examine the localization and function of CG31030, a novel ATP6AP1 homolog in *Drosophila* whose unique characteristic is to be expressed selectively and ubiquitously in neurons. Whereas a complete deficiency of this protein is lethal, we found that partial *CG31030* knockdown in larval motoneurons impaired synaptic vesicle acidification, reduced quantal size, which is the amplitude of the postsynaptic response to the release of a single synaptic vesicle, and induced severe locomotion defects. We also report that CG31030 from brain tissue co-immunoprecipitated with V-ATPase subunits of the V_0_ domain. Overall, our results indicate that CG31030 is a novel accessory subunit of the neuronal V-ATPase that appears to be involved in the regulation of synaptic activity.

## Results

### CG31030 is specifically expressed in neurons and addressed to synaptic areas

Here we studied in *Drosophila* the gene *CG31030*, identified in FlyBase as a paralog of *VhaAC45*/ *CG8029*, and as an ortholog of vertebrate *ATP6AP1*/*AC45* (Thurmond et al., 2019). CG31030 and VhaAC45 indeed share 69.9% similarity in amino-acid sequences (Supplementary Figure 1A). *CG31030* is also classified in the V-ATPase family group by the InterPro database (accession: Q8IMJ0_DROME) (Mitchell et al., 2019). According to FlyAtlas (Chintapalli et al., 2007), *VhaAC45* is expressed ubiquitously in *Drosophila* tissues. In contrast and interestingly, *CG31030* seems to be specifically expressed in the nervous system in both larval and adult flies, making it a possible candidate for the specific regulation of neuronal V-ATPase (Supplementary Figure 1B). To confirm this prediction, we checked by RT-qPCR the repartition of *CG31030* transcripts in three parts of the adult fly body: the head and thorax, which contains the brain and ventral nerve cord (VNC), respectively, and the abdomen, which is relatively poor in nervous tissue. Both females and males showed highest expression in the head, minor expression in the thorax and no detectable expression in the abdomen (Figure 1A). The expression of *CG31030* therefore closely follows the repartition of the nervous system.

**Figure 1.**
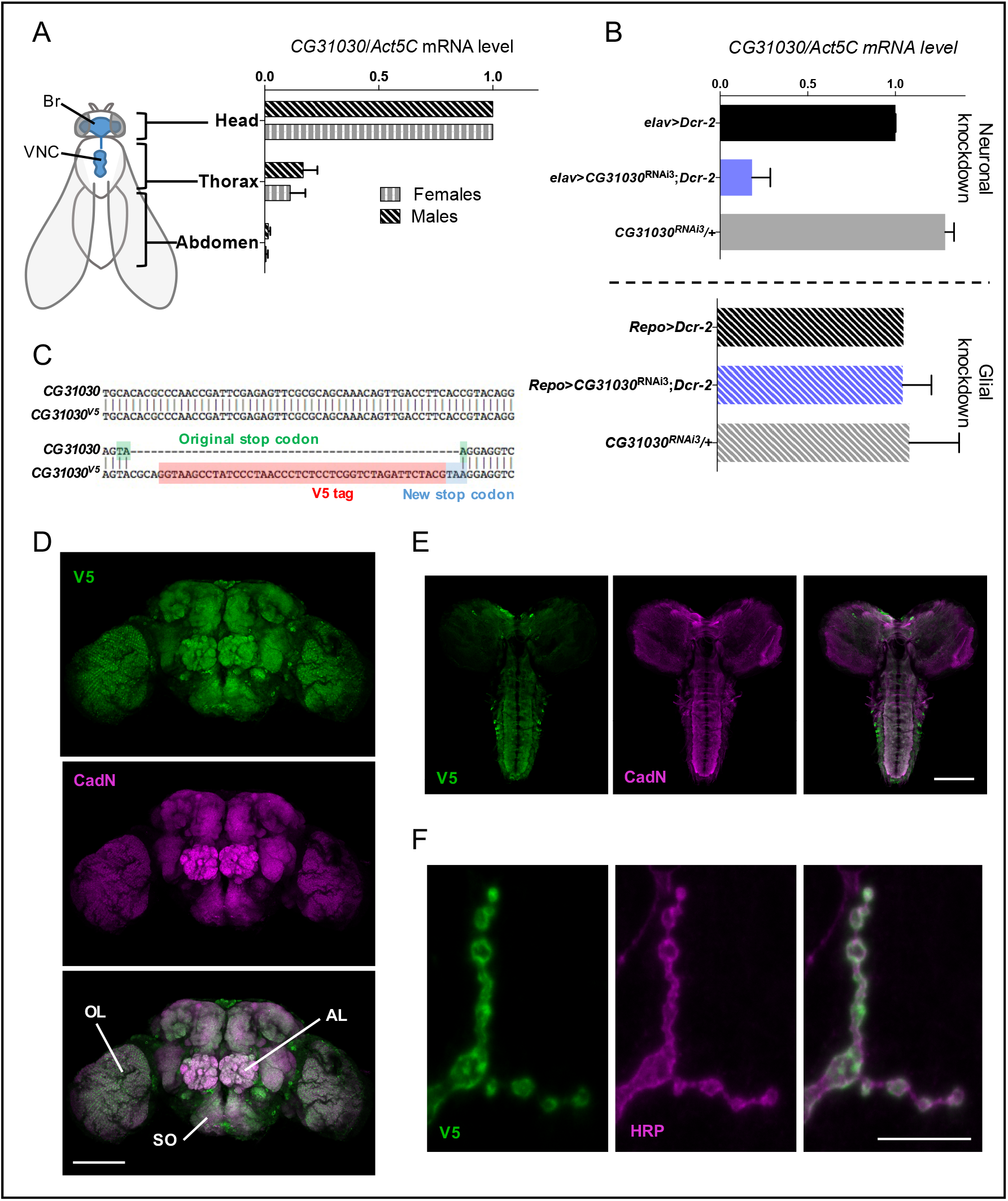
CG31030 is expressed in neurons and addressed to synaptic areas. (**A**) In both males and females, *CG31030* mRNA abundance follows the localization of the CNS (shown in blue on the fly sketch), with the highest expression in the head. Br: brain, VNC: ventral nerve cord. Results of three independent experiments. (**B**) Expression of *CG31030*^RNAi3^ with *Dcr-2* in all neurons using *elav-Gal4* decreased *CG31030* mRNA level in head by more than 80%, while expression of the RNAi construct and Dcr-2 in all adult glial cells with *repo-Gal4* had not effect. Results of three independent experiments. Mean values with SD are reported on the graphs. (**C**) Construction of the *CG31030*^V5^ mutant strain. A V5 tag (in red) was fused to the C-terminal end of the CG31030 protein by inserting the V5 coding sequence ended by a new stop codon (in blue) in place of the original stop codon (in green) in the *CG31030* gene using the CRISPR-Cas9 technology. (**D, E**) Anti-V5 immunostaining in the *CG31030*^V5^ strain revealed that CG31030 is mainly addressed to synaptic areas in the adult brain (**D**) and larval CNS (**E**), as indicated by its apparent colocalization with the presynaptic marker Cadherin-N (CadN). OL: optic lobe, AL: antennal lobe, SO: subœsophageal ganglion. (**F**) A V5-immunopositive signal was also detected at the neuromuscular junction of *CG31030*^V5^ larvae, and found to co-localize with anti-horseradish peroxidase (HRP) immunostaining that labels neuronal membranes, confirming the synaptic localization of *CG31030*. Scale bars: 100 μm in D and E, and 10 µm in F.

The single-cell RNA-Seq Scope database (Davie et al., 2018) furthermore indicated that, in *Drosophila*, CG31030 is expressed in all neurons, with few or no expression in glial cells. To verify this, we expressed three different *CG31030* RNAi either with either the pan-neuronal driver *elav-Gal4* or the pan-glial driver *repo-Gal4* (see Supplementary Table 3 for genotypes of the different *Drosophila* lines used in this study). The expression of *CG31030*^RNAi1^ and *CG31030*^RNAi2^ was found to be lethal at embryonic and 1^st^ larval stages, respectively, while *CG31030*^RNAi3^ produced viable adults with a shortened longevity (Supplementary Figure 2), and obvious locomotor impairments. This difference in phenotypes observed with different RNAi constructs could be attributed to a variation in residual levels of the CG31030 protein. RT-qPCR experiments showed that the pan-neuronal expression of *CG31030*^RNAi3^ together with the RNAi booster *Dicer-2* (*Dcr-2*) was sufficient to decrease by more than 80% *CG31030* transcripts abundance in extracts from the adult heads (Figure 1B). On the other hand, glial expression of this RNAi construct with *repo-gal4* had no significant effects on *CG31030* transcript level (Figure 1B), confirming that this gene is selectively expressed in neurons. It is interesting to note that both *CG31030*^RNAi1^ and *CG31030*^RNAi2^ also induced lethal phenotypes at various developmental stages when expressed either with a glutamatergic (*VGlut-Gal4*) or a cholinergic (*Cha-Gal4*) neuronal driver (data not shown), in accordance with the RNA-Seq data of the Scope website suggesting a pan-neuronal expression of this gene.

To validate these observations, we placed the available MIMIC line *CG31030*^MI107^ that contains a stop codon inserted in the middle of the gene (Nagarkar-Jaiswal et al., 2015) over the deficiency *Df(3R)Exel6214* encompassing *CG31030* (Parks et al., 2004). The resulting mutant, likely to be a null, was found to be embryonic lethal, in agreement with the results obtained with two *CG31030* RNAi lines. Remarkably, re-expressing the gene selectively in neurons, using a *UAS-CG31030* construct driven by *elav-Gal4*, was sufficient to rescue this lethality, producing viable and fertile adults with no obvious behavioral defects (Supplementary Table 1). These results strongly indicate that *CG31030* is an essential gene whose expression appears to be specifically required in neurons.

Next, we studied the cellular localization of this protein. We used the CRISPR-Cas9 technique to insert a small V5 epitope tag in frame at the 3’ end of its gene, disrupting the stop codon, thus generating the *CG31030*^V5^ mutant line (Figure 1C). By immunostaining with a V5-specific antibody, we observed that the general expression pattern of CG31030 in adult brain (Figure 1D) and larval CNS (Figure 1E) was widespread and quite similar to that of the synaptic marker Cadherin-N (CadN), indicative of a predominantly synaptic localization. No specific signal was detected in a control *w*^1118^ line that does not contain the V5-tagged protein (not shown). Some neuronal cell bodies were also marked with the V5 antibody in the *CG31030*^V5^ line, both in adult and larva, and the synapse-containing neuropile areas (like the antennal lobes) were not as sharply defined as with the CadN antibody, suggesting that axons could also be immunopositive. Finally, co-immunostaining of larval body muscles wall with anti-horseradish peroxidase (HRP) antibodies, a marker of *Drosophila* neurons (Jan and Jan, 1982), showed precise co-localization with the V5 signal, indicating that CG31030 is addressed to synaptic boutons at the larval neuromuscular junction (Figure 1F).

### CG31030 co-immunoprecipitates with V-ATPase proteins

The CG31030 protein is predicted to be part of the InterPro V-ATPase family, but no experimental information is currently available regarding its potential interactors. To determine whether CG31030 could interact, directly or indirectly, with subunits of the V-ATPase complex, we carried out co-immunoprecipitation experiments using an anti-V5 antibody on proteins extracted from heads of *CG31030*^V5^ mutants and *w*^1118^ control flies, followed by nano LC-MS/MS mass-spectrometry analysis of the precipitated proteins. Three independent experiments were performed to increase reliability, and in total, 410 proteins were identified in all three experiments. Among those, only 12 proteins had at least a two-fold abundance difference with the control in all three experiments (Figure 2A and Supplementary Table 2), one of them being, as expected, the co-immunoprecipitation target CG31030. Remarkably, three of the 11 other proteins were identified as subunits of the V-ATPase complex: Vha100-1, VhaAC39-1 and ATP6AP2 (Figure 2B).

**Figure 2.**
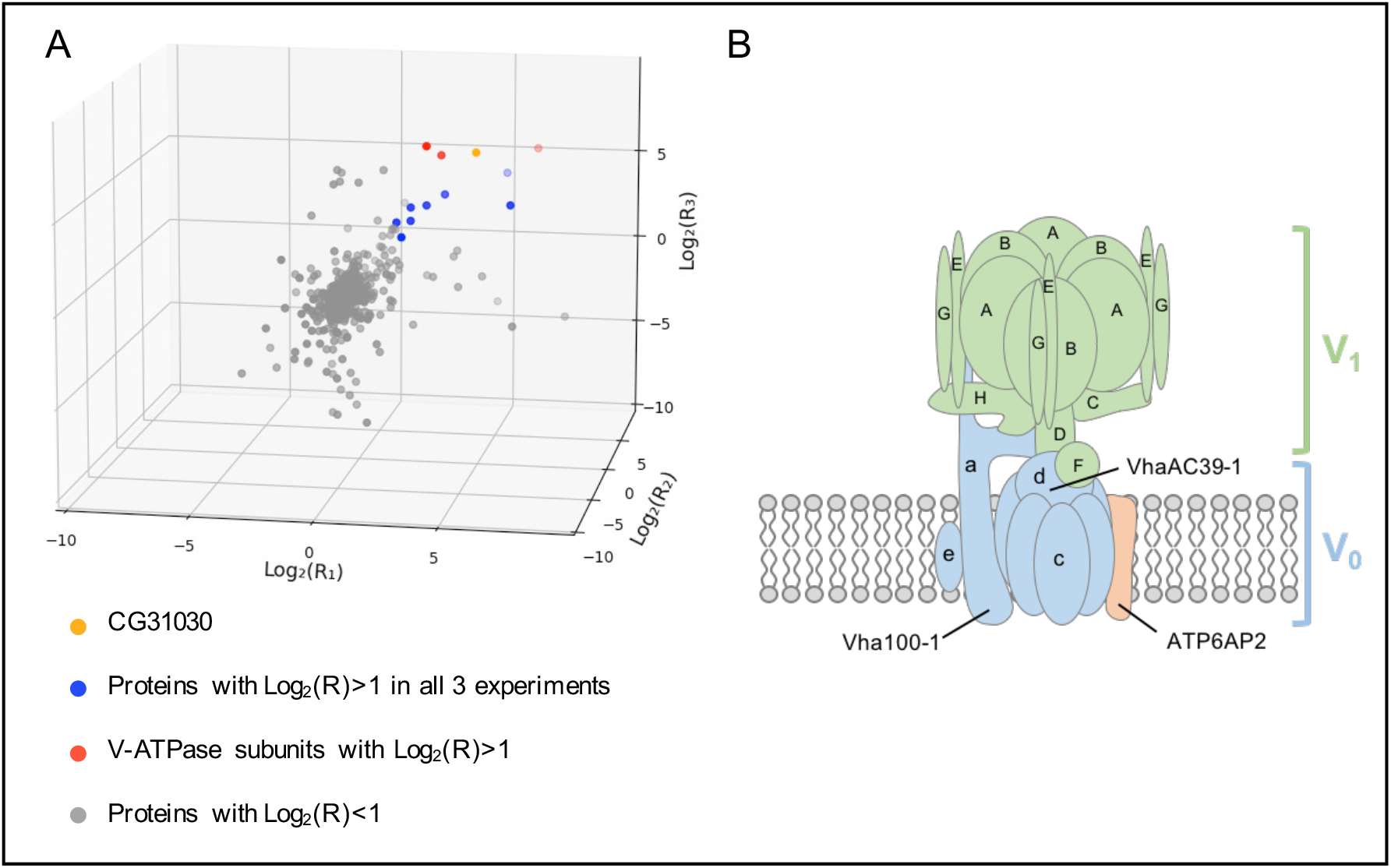
CG31030 co-immunoprecipitates with V-ATPase subunits. (**A**) Scatter plot of proteins identified by nano LC-MS/MS in three independent co-immunoprecipitation experiments using anti-V5 antibodies. R1 and R2 represent the abundance ratio of proteins identified in adult head extracts from *CG31030*^V5^ over *w*^1118^ control in experiments 1 and 2, respectively. Solid lines indicate Log2(R) = 1, which corresponds to a two-fold abundance difference. 12 proteins were found to be at least twice as abundant in *CG31030*^V5^ as in the control in all three experiments (red or blue dots on the graph). One of them is the immunoprecipitation target CG31030 (yellow dot) and three of these proteins belong to the V-ATPase complex (red dots). A list of these 12 proteins with their Log2(R) values is provided in Supplementary Table 2. (**B**) Standard model of the *Drosophila* V-ATPase complex showing structure of the V1 and V0 domains and the predicted localization of the three subunits that co-immunoprecipitated with CG31030.

*Vha100-1* and *VhaAC39-1* code for subunits *a* and *d* of the V_0_ domain, respectively (Vasanthakumar and Rubinstein, 2020). The subunit *a*, coded by five different genes in *Drosophila*, is the proton port of the pump (Collins and Forgac, 2020). Among the five isoforms, Vha100-1, which co-immunoprecipitated with CG31030 in our experiments, has been shown to be specifically required in neurons and present at the synapse (Hiesenger et al., 2005). Subunit *d* of V_0_ is coded by two *Drosophila* genes: *VhaAC39-1* and *VhaAC39-2*, and only the first co-immunoprecipitated with CG31030. According to FlyAtlas, VhaAC39-1 is expressed in many tissues and enriched in the brain, while VhaAC39-2 seems to be mostly found in testis and salivary glands. For both V_0_ subunits, CG31030 thus co-precipitated with the likely neuronal isoform. The co-immunoprecipitated V-ATPase subunit which appeared to be the most enriched in the *CG31030*^V5^ sample was interestingly the accessory subunit ATP6AP2, suggesting a possible direct interaction between this protein and CG31030 (Supplementary Table 2). These experiments therefore reinforce the hypothesis that CG31030 directly interacts with the neuronal V-ATPase complex, and more specifically with V_0_ since all detected partners belong, or interact, with this domain.

### CG31030 knockdown increases the pH of synaptic vesicles

Because CG31030 appeared to be mainly localized in synaptic areas (see Figure 1), we chose to look at the physiological effect of its disruption at the *Drosophila* larval neuromuscular junction, a model that has contributed to the study of many essential synaptic processes. At synaptic nerve endings, a prominent role of the V-ATPase is to acidify the lumen of synaptic vesicles, the electrochemical gradient generated providing the driving force to load and concentrate the neurotransmitters. Thus, a malfunction of synaptic V-ATPase should induce a decrease of neurotransmitter concentration inside the vesicles, potentially resulting in an altered synaptic transmission. To test this hypothesis, we co-expressed each of the two strongest *CG31030* RNAi constructs together with VMAT-pHluorin, a pH-sensitive probe targeted to synaptic vesicles (Wu et al., 2013), in larval motoneurons using the glutamatergic driver *OK371-Gal4.* Both RNAi1 and RNAi2 induced a lethal phenotype at pupal stage in these conditions. VMAT-pHluorin is an ecliptic pHluorin that is fluorescent at neutral pH, and gets quenched, by protonation, at acidic pH (Miesenböck et al., 1998). Thus, in control condition, VMAT-pHluorin should not be fluorescent in synaptic vesicles, whose pH is at around 5.5, but only when externalized on the presynaptic membrane during exocytosis, and so, in contact with the more neutral synaptic cleft milieu. In the case of an acidification defect of synaptic vesicles, the probe could be fluorescent both in synaptic vesicles, where pH would be abnormally high, and on the presynaptic membrane (Figure 3A, central panel).

**Figure 3.**
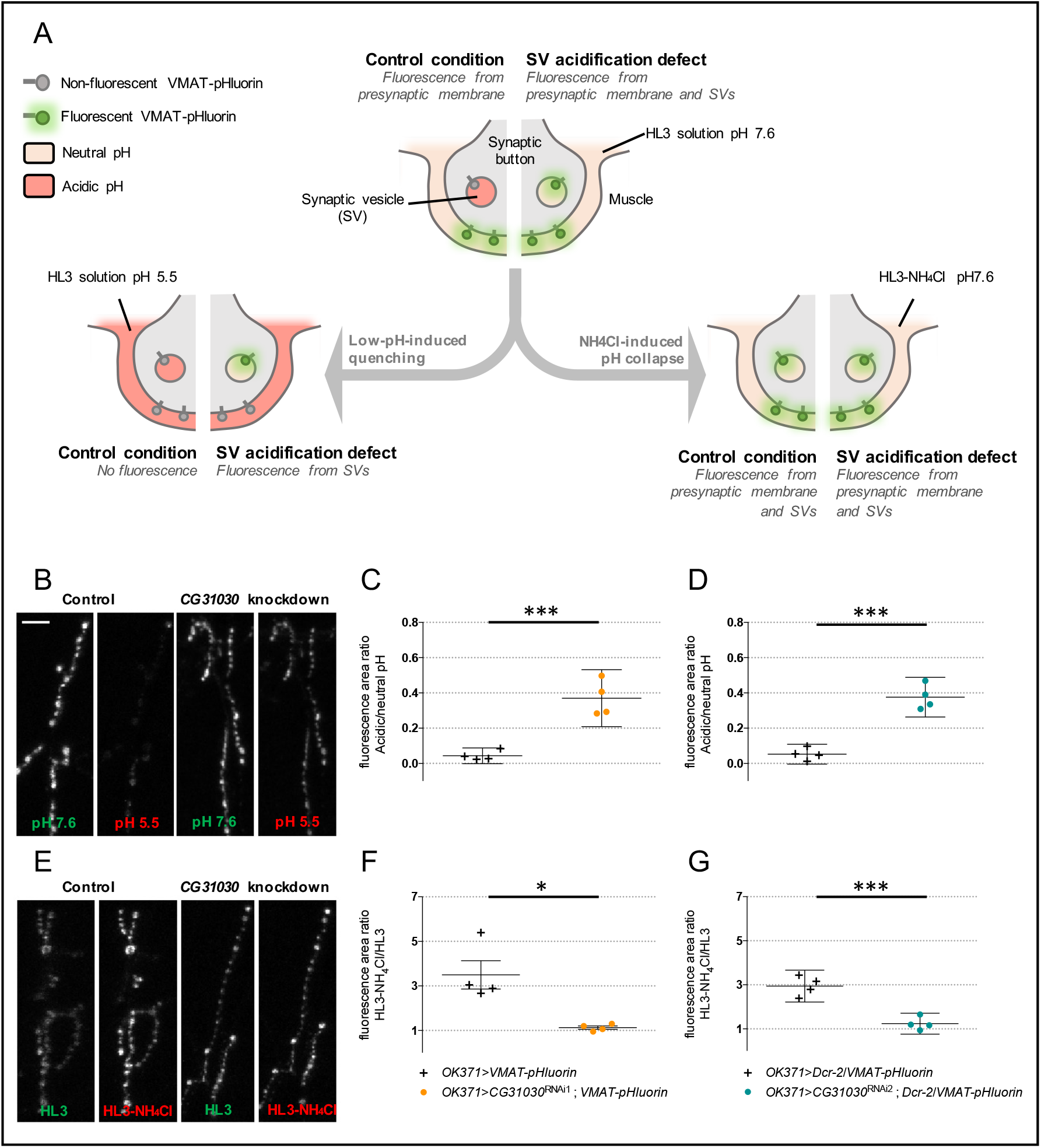
CG31030 knockdown larvae have a synaptic vesicle acidification defect. (**A**) Schematic representation of the protocols used to assess relative acidity levels of synaptic vesicles at the larval neuromuscular junction. (*Top center diagram*) In control conditions, fluorescence can be emitted by VMAT-pHluorin in the presynaptic membrane but not in synaptic vesicles since their lumen is acidified. In case of defective synaptic vesicle acidification, both the presynaptic membrane and synaptic vesicles should emit fluorescence. (*Left diagram*) The fluorescence emitted by VMAT-pHluorin in the presynaptic membrane can be quenched by replacing the extracellular medium with an acidic HL3 solution. This quenching should result in an almost complete extinction of the signal in control flies, in which synaptic vesicles are normally acidified, while a residual signal is expected to be visible in flies having a synaptic vesicle acidification defect. (*Right diagram*) Replacement of 50 mM NaCl by 50 mM NH4Cl in the neutral HL3 solution should lead to a collapse of the pH gradients due to the free diffusion of NH3 in membranes, so that fluorescence will be emitted both by the presynaptic membrane and synaptic vesicles both in control and mutant conditions. (**B**) Representative pictures showing the effect of perfusing an acidic HL3 solution on VMAT-pHluorin fluorescence in control and *CG31030* knockdown larvae. (**C**, **D**) Quantification of the ratio of the fluorescence level at pH 5.5 over the original signal at pH 7.6. Whereas the low pH extinguished fluorescence in control flies, about 37% of the signal persisted after quenching in *CG31030* knockdown larvae using two different RNAi constructs. (**E**) Representative pictures showing the effect of collapsing the synaptic vesicle pH by perfusing HL3-NH4Cl in control and *CG31030* knockdown larvae. (**F**, **G**) Quantification of the ratio of the signal in HL3-NH4Cl over the original signal in HL3 showed that fluorescence increased about 3-fold in controls while it only rose by 10 to 20% depending on the RNAi in *CG31030* knockdown larvae. Results of four independent experiments, with 3-5 larvae analyzed per genotype in each experiment. Unpaired Student’s *t*-test, \**p* < 0.05 \*\**p* < 0.01. Mean values with 95% confidence intervals are reported on the graphs. Scale bar: 15 μm in B and E.

In order to evaluate the ratio of the internal fluorescence (from synaptic vesicles) over the external fluorescence (from the presynaptic membrane) at the neuromuscular junction of *CG31030* knockdown larvae compared to controls, we first quenched the external signal by replacing the physiological milieu by an identical one with pH adjusted to 5.5 (Figure 3A, left panel). This operation resulted in only the internal signal being conserved. In controls, this meant that all signal was abolished, as expected because the synaptic vesicles were normally acidified (Figure 3B, left panel). In contrast and strikingly, a residual signal was still visible in this acidic milieu in both RNAi1 and RNAi2 knockdown larvae (Figure 3B, right panel). Quantification of the ratio of fluorescence area in acidic milieu over neutral milieu showed that about 37% of the total signal remained visible in the RNAi larvae after external quenching (Figure 3C and D). To verify that the residual signal seen in knockdown larvae was indeed coming from inside vesicles, the opposite strategy was used: instead of quenching the outside signal, we revealed all the internal one by collapsing the pH gradient of synaptic vesicles (Figure 3A, right panel). To do so, we replaced the physiological milieu by an ammonium solution, as previously described (Poskanzer and Davis, 2004). This solution had the same composition except that 50 mM NaCl were replaced by 50 mM NH_4_Cl. Ammonium and ammonia being in equilibrium (NH_4_^+^ ⇄ NH_3_ + H^+^), uncharged ammonia crosses membranes and binds to protons leading to an alkalization of vesicle lumen pH. This increase of vesicular pH is maintained during NH_4_Cl exposure. In control condition, this gradient collapse should reveal the VMAT-pHluorin probe present in synaptic vesicles and thus highly increase the fluorescent signal. On the contrary, in the case of an acidification defect, the signal should remain fairly stable since synaptic vesicles are already fluorescent. Results were consistent with this hypothesis: while the fluorescence of controls increased about 3 folds in the pH collapsing ammonium solution, the signal in *CG31030* knockdown larvae hardly rose by 10 to 20% depending on the RNAi construct (Figure 3E, F and G). Taken together, these results indicate that *CG31030* knockdown significantly decreases protons concentration in synaptic vesicles of motoneuron terminals.

### CG31030 downregulation decreases larval locomotor performance

A reduced pH gradient of synaptic vesicles in larval motoneurons could alter synaptic transmission and, consequently, the larval locomotor behavior. To assess locomotion, we recorded the spontaneous crawling of 3^rd^-instar larvae expressing *CG31030* RNAi1 or RNAi2 in motoneurons with *OK371-Gal4* on an agar plate with no food source for 2-min periods. Tracking was then performed using the FIMtrack software (Risse et al., 2014), allowing measurement of the total distance travelled and of the stride size, defined as the distance crawled during one peristaltic wave of muscle contraction, and duration (Figure 4A). The knockdown larvae obviously moved less than controls on the plate, as confirmed by their trajectory maps (Figure 4B and C), travelling a distance only half as long as controls with both RNAi (Figure 4E and F. This effect was not simply due to a difference in larval size, as the average length and width of recorded larvae were not significantly different (Supplementary Figure 3).

**Figure 4.**
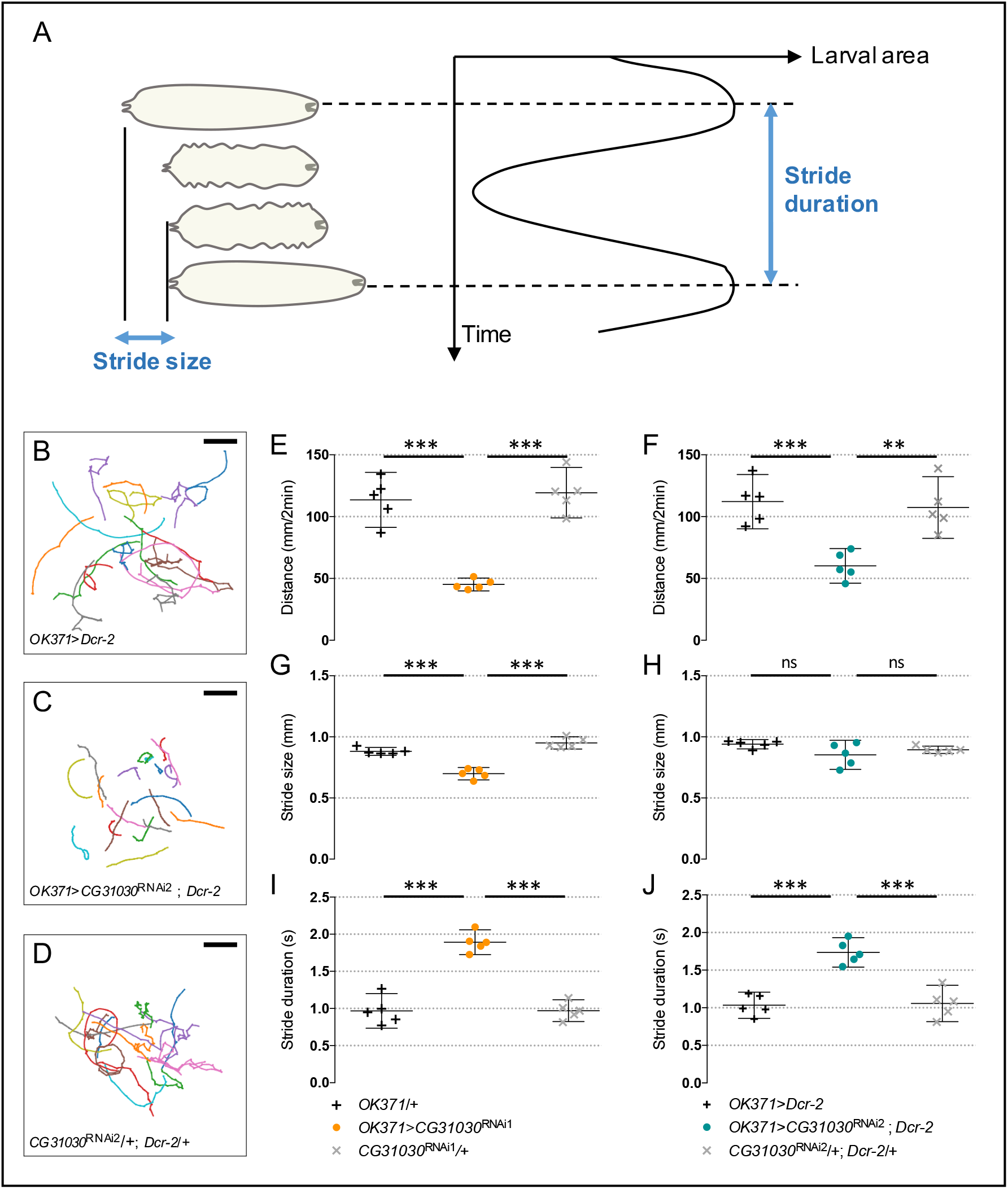
CG31030 downregulation in motoneurons decreases larval locomotor performance. (**A**) Schematic representation of the successive phases of larval locomotion. Stride size is defined as the distance crawled during one peristaltic wave of muscle contraction, while stride duration is the time necessary for the completion of one peristaltic wave. (**B**-**D**) Locomotor trails of individual larvae recorded over a period of 2 min. Larvae expressing *CG31030* RNAi in motoneurons show reduced spontaneous movements (**C**) compared to the driver and UAS controls (**B** and **D**, respectively). Scale bar: 25 mm. (**E**, **F**) Quantification of the travelled distances confirmed that knocking down *CG31030* in motoneurons with two different RNAi induced significant locomotor defects. (**G**-**J**) Stride size (**G**, **H**) appears to be much less affected than stride duration (**I**, **J**) in the knocked-down larvae. Result of five independent experiments, with 3-4 larvae analyzed per genotype in each experiment. One-way ANOVA with Dunnett’s post-test for multiple comparisons, \*\**p* < 0.01, \*\*\**p* < 0,001. Mean values with 95% confidence intervals are reported on the graphs.

In contrast, the stride size actually showed little or no difference, depending on the RNAi, between knockdown larvae and controls (Figure 4G and H). Instead, we found that the stride duration, which is the time necessary to accomplish one peristaltic wave, was significantly longer in knockdown animals compared to controls for both RNAi (Figure 4I and J). These results suggest that the knockdown larvae may need about twice as much time as controls to reach the required level of muscular contraction to accomplish one stride. This could be a compensatory mechanism to adapt to a lower amount of neurotransmitter released in response to motor nerve stimulation, as would be expected if synaptic vesicles are less filled with neurotransmitter in *CG31030* knockdown context.

### Quantal size is reduced in *CG31030* knockdown larvae

The locomotor deficit of *CG31030* knockdown larvae could originate from an impairment of synaptic transmission at the neuromuscular junction or a defect in the central control of motor behavior, or both. To determine if neurotransmission was affected, we carried out electrophysiological recordings to measure the quantal size, which is the postsynaptic response to the release of one synaptic vesicle, at the larval neuromuscular junction. We expressed *CG31030* RNAi1 or RNAi2 in motoneurons with the *OK371-Gal4* driver, and recorded spontaneous miniature excitatory postsynaptic potentials (mEPSPs) intracellularly from ventral longitudinal abdominal muscle 6 of segment A3 (Figure 5A). These muscles are innervated by synaptic boutons that are clearly marked by *OK371-Gal4*, as shown by the co-localization of membrane-associated GFP and the postsynaptic marker Discs large (Dlg) in *OK371*>*mCD8::GFP* flies (Figure 5B). Representative amplitude distributions of mEPSPs in a control and RNAi1 larvae are shown in Figure 5C. Quantal analysis of recorded events confirmed that both RNAi1 and RNAi2 knockdown larvae have a significantly reduced quantal size compared to controls (Figure 5D), suggesting a decrease in glutamate vesicular uptake, and potentially linking vesicle acidification defect and locomotor impairment. The V-ATPase dysfunction in synapses of *CG31030* knockdown larvae apparently led to incomplete loading of vesicles, thus decreasing the amount of neurotransmitter released per unit of time during a peristaltic wave, and potentially slowing down larval locomotion. Although the mean frequency of mEPSPs of both RNAi1 and RNAi2 larvae appeared lower than controls, this effect was not statistically significant (Figure 5E). This result suggests that *CG31030* knockdown did not significantly increased the number of unacidified vesicles empty of neurotransmitter.

**Figure 5.**
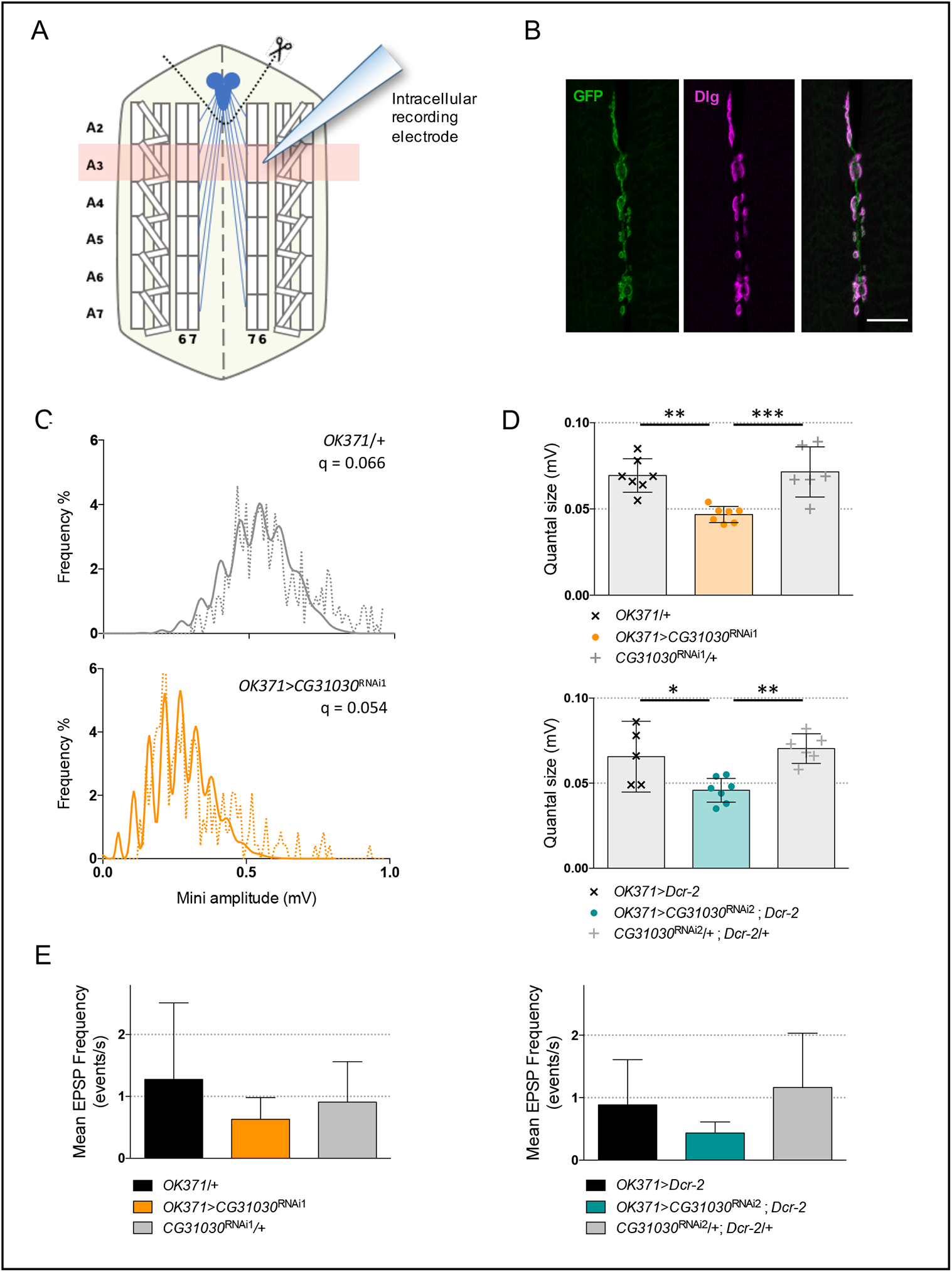
Synaptic quanta size is reduced in *CG31030* knockdown larvae. (**A**) Schematic representation of a dissected larval fillet. Spontaneous excitatory postsynaptic potentials (EPSPs) were recorded intracellularly from the ventral longitudinal abdominal muscle 6 in segment A3. (**B**) Expression of membrane-associated mCD8::GFP with the glutamatergic driver *OK371-Gal4* strongly labels the presynaptic nerve endings at the neuromuscular junction of muscles 6-7 in segment A3. The scaffolding protein Discs large (Dlg) was used as a postsynaptic marker. Scale bar: 20 μm. (**C**) Representative distributions of spontaneous mEPSPs recorded in a control larva (*top panel*, *in grey*) and a *CG31030* RNAi knockdown larva (*bottom panel, in orange*). The dotted lines represent actually recorded amplitudes and the plain lines the computed theoretical distributions. Genotypes and quantal size (q) are indicated on each graph. (**D**) Quantal analysis of recorded events showed that knockdown larvae have a significantly reduced synaptic quanta size compared to controls, both with *CG31030* RNAi1 (*top panel, in orange)* or RNAi2 (*bottom panel*, *in blue*). (E) Although both RNAi1 (*left panel*) and RNAi2 (*right panel*) larvae had a lower mean EPSP frequency than controls, this difference was not statistically significant. One-way ANOVA with Dunnett’s post-test for multiple comparisons, \**p* < 0.05, \*\**p* < 0.01, \*\*\**p* < 0.001. Mean values with 95% confidence intervals are reported on the graphs.

## Discussion

In this study, we investigated the hypothesis that the previously uncharacterized *Drosophila* protein CG31030 is a specific regulator of the neuronal V-ATPase. At variance with its broadly expressed paralog VhaAC45, we have shown that CG31030 is found mainly, if not only, in neurons. We also provide evidence that CG31030 interacts with two constitutive subunits and one accessory subunit of the V-ATPase, the constitutive ones being also enriched in neurons, and that it is required to have properly acidified synaptic vesicles. This implies that CG31030 is an essential protein for nervous system functioning in *Drosophila*.

### CG31030 is an essential synaptic protein

In yeast, all V-ATPase subunits are coded by a single gene, with the exception of the V_0_ subunit *a*. The knock-out of any of the single-gene subunits all present a similar phenotype: the inability to survive in a neutral pH environment (Nelson, 2003). For subunit *a*, the same phenotype was only achieved in a double-mutant of both isoforms (Manolson et al., 1994). In multicellular organisms, mutations of V-ATPase subunits, or accessory proteins, also often lead to a lethal phenotype at various developmental stages, whether in mice (Inoue et al., 1999; Sun-Wada et al., 2000; Schoonderwoert et al., 2002-b), *C. elegans* (Lee et al., 2010), or flies (Davies et al., 1996; Allan et al., 2005). Similarly, a *CG31030* null mutant was found to be embryonic lethal, and, interestingly, this lethality could be rescued to the adult stage by re-expressing *CG31030* in all neurons, showing that the protein is specifically required in this cell type for fly survival. The percentage of adult survivors was about three times less than what would be expected in case of a full rescue (Supplementary Table 1) and the missing two-thirds likely died at an early developmental stage as no larval lethality was observed. This could be explained by the fact that CG31030 protein re-expression in rescued knock-out flies first required the expression of Gal4 under regulation of the *elav* promoter, which starts to express rather late in embryos, followed by activation of the UAS sequence upstream the *CG31030* insert. So, it is possible that part of the mutant embryos did not survive the delay inherent to this ectopic expression process. We found that *CG31030* transcripts follows the repartition of the nervous system, being mainly expressed in the head. Moreover, pan-neuronal expression of *CG31030*^RNAi3^, the RNAi construct with the weakest effect on fly survival, was sufficient to decrease by more than 80% its level in the brain of adult escapers. These results indicate that CG31030 expression is mainly neuronal, if not even entirely restricted to neurons. In addition, CG31030 cellular localization shows similarity with a synaptic pattern. Cell bodies were also marked, although generally less strongly. Synaptic protein complexes can be assembled in the cell bodies before being transported in axons, and it is the case for the soluble and membrane-bound domains of synaptic V-ATPase, as was shown in *Torpedo* (Morel et al., 1998). It is difficult to decide whether the signal coming from cell bodies corresponds to a functional site for CG31030 or rather to newly produced proteins that have not yet been targeted to synapses. However, the prominent signal in synaptic areas led us to assume that CG31030 plays an important role in the synaptic process.

### CG31030 is required for synaptic vesicle acidification

In presynaptic terminals, the most abundant organelles are synaptic vesicles, which require acidification to provide the driving force for neurotransmitter loading. This acidification is ensured by the V-ATPase pump, which generates an electrochemical gradient by importing protons into the vesicle lumen, thus creating both a membrane potential (ΔΨ, inside positive voltage) and a pH gradient (ΔpH, acidic lumen). Our experiments showed that the knockdown of *CG31030* at the glutamatergic larval neuromuscular junction increased the internal pH of synaptic vesicles, and so decreased the ΔpH component of the electrochemical gradient generated by the V-ATPase. This result is however by itself insufficient to conclude to a dysfunction of the proton pump, since other players have been shown to influence synaptic vesicle pH gradient downstream V-ATPase activity. For example, cation/H^+^ exchangers, found on synaptic vesicles, can decrease ΔpH while increasing ΔΨ by exchanging cations, like Na^+^ or K^+^, against luminal H^+^ (Takamori, 2016). This activity can be upregulated by increased intracellular concentration of these cations (Huang and Trussell, 2014). Glutamate being negatively charged, its transport across vesicular membrane is predominantly driven by ΔΨ (Maycox et al., 1988; Tabb at al., 1992). As a consequence, the increased membrane potential resulting from up-regulation of cation/H+ exchangers actually facilitates glutamate uptake (Goh et al., 2011; Huang and Trussell, 2014). We measured synaptic quanta size by quantal analysis of mEPSPs of *CG31030* knockdown larvae and found that it was decreased, contrary to what would be expected in the case of an upregulation of cation/H^+^ exchanger activity. This apparent decrease of vesicular glutamate concentration was corroborated by its phenotypical consequence, namely the locomotion defect exhibited by knockdown larvae. This suggests that the observed diminution of ΔpH is correlated with a diminution of ΔΨ, thus pointing towards a V-ATPase malfunction. This hypothesis is further supported by the co-immunoprecipitation of CG31030 with three V-ATPase subunits of the V_0_ domain (Vha100-1, VhaAC39-1 and ATP6AP2), all of them being associated with the neuronal V-ATPase. Altogether, these results strongly suggest that CG31030 is a neuronal protein necessary to V-ATPase function in synapses.

Some V-ATPase subunits isoforms in other species have been shown to target specific subcellular compartments, like the *Torpedo* V_0_ a1 isoform which is found in synaptic V-ATPase and not in neuronal cell bodies (Morel et al., 2003). Similarly, CG31030 could be specifically involved in the regulation of synaptic V-ATPase, or, alternatively, globally act on all neuronal V-ATPase. Specificity level could even be pushed further, since synaptic vesicles are not the only organelles requiring acidification that are localized in synapses. Thus, a synaptic V-ATPase regulator could be devoted to synaptic vesicles as it could be affecting more generally all synaptic V-ATPase complexes, indifferently of their membrane localization. The answer to such questions could help better understand regulations of synaptic transmission, since synaptic V-ATPase activity is one of the pre-synaptic modulators of quantal response (Takamori, 2016; Gowrisankaran and Milosevic, 2020).

### CG31030 interacts with ATP6AP2 and may regulate V-ATPase domain dynamics

While accessory subunits directly interact with V-ATPase domains, other stimulus can indirectly affect activity of the complex, like glucose concentration or serotonin (Sautin et al., 2005; Voss et al., 2007). Co-precipitation of CG31030 with V-ATPase subunits does not necessarily imply a direct interaction, but suggests *a minima* an indirect association with the complex, possibly in a non-transient manner. The impairment in V-ATPase activity induced by CG31030 downregulation rules out the hypothesis of an inhibitory action of this protein on the proton pump. Nevertheless, the precise requirement of CG31030 for activity of the V-ATPase complex still remains unknown to date. Lethality of the mutant could indicate an essential role of CG31030 in V-ATPase function. However, some V-ATPase regulators and accessory subunits have been found in other signaling pathways, like ATP6AP2 in the renin-angiotensin system (Nguyen, 2010), so we cannot exclude that lethality could be due to CG31030 playing a part in other vital neuronal functions. High signal in axons could also point to a role in targeting the complex to the synapses. The two domains of the V-ATPase, V0 and V1, are believed to be assembled in cellular bodies before being separately transported to synaptic area, V_0_ by fast axonal anterograde transport, most likely directly on new synaptic vesicles, and V_1_ by slow axonal transport like other cytoplasmic synaptic proteins (Morel et al., 1998). CG31030 could be involved in the transport of one of the two domains, and effects of CG31030 knockdown could then result from a decreased synaptic abundance of the affected V-ATPase domain. However, it is believed that only one copy of the V-ATPase complex is sufficient to properly acidify one synaptic vesicle (Takamori et al., 2006), a consequence of this being that neurotransmitter loading would be an all-or-none process. Consistent with this, mutation of the *Drosophila* synaptic V_0_ subunit Vha100-1 does not seem to impact quantal size but rather mEPSP frequency, potentially reflecting the presence of an increased number of empty synaptic vesicles (Hiesinger et al., 2005). Similarly, a reduction in the frequency of spontaneous quantal events with no change in quantal size was observed when the *Drosophila* vesicular glutamatergic transporter (VGlut) was downregulated, also suggesting that a single copy of VGlut is sufficient for proper acidification of a vesicle (Daniels et al., 2006). The fact that CG31030 knockdown changes the quantal size but not significantly mEPSP frequency seems to point toward a role of this accessory protein in V-ATPase efficiency, rather than in a process affecting the abundance of the complex such as synaptic targeting.

In this respect, it is interesting to note that the V-ATPase protein that more consistently co-immunoprecipitated with CG31030 in our experiments is the accessory subunit ATP6AP2, the fly homolog of human ATP6AP2/PRR. Interestingly, the vertebrate homolog of CG31030, ATP6AP1/Ac45, has also been shown to interact with ATP6AP2, and the complex they form has been proposed to enable the assembly and disassembly of the catalytic and membrane domains of the V-ATPase in the mammalian brain (Rujano et al., 2017; Abbas et al., 2020). This suggests that CG31030 could similarly play a role in the regulation of these dynamic processes, which may also influence neurotransmitter loading and release in a more continuous way. Our results suggest indeed that modulating V-ATPase activity in presynaptic terminals can finely affect quantal size. Whether such regulation actually occurs in physiological, or pathological, conditions remains to be established. Owing to its structural and functional similarity with its closest *Drosophila* homolog, VhaAC45/ATP6AP1, we propose therefore to name CG31030 “VhaAC45-Like” (VhaAC45L) in further work.

Human *ATP6AP2*/*PRR* is known to be a Parkinsonism candidate gene and its mutations in humans, mice or flies can lead to cognitive impairment, neurodegeneration and epilepsy (Korvatska et al., 2013; Dubos et al. 2015; Ichihara and Yatabe, 2019). It is therefore of major interest to identify a potential new interactor of ATP6AP2 that is specific to the nervous system, as it could help better understand the consequences of V-ATPase dysregulation on synaptic transmission in pathological contexts. Accordingly, it would be very interesting to determine whether human ATP6AP1 and/or ATP6AP1L share a conserved function with *Drosophila* CG31030/VhaAC45L in the nervous system. Every new actor identified allows us to paint a more detailed picture of the complex regulations surrounding neuronal V-ATPase specificity, providing new angles for potential therapeutic targets, and a better understanding of fundamental processes such as synaptic transmission.

## Materials and methods

### *Drosophila* strains, construction and culture

Flies were raised on standard agar-cornmeal-yeast medium, at 25°C in a 12:12h light-dark cycle. *CG31030* RNAi strains and mutants were obtained from the Bloomington *Drosophila* stock center (BDSC) and the Vienna Drosophila Resource Center (VDRC). Detailed genotypes and references of these lines are provided in Supplementary Table 3. To construct the *UAS-CG31030* strain, the *CG31030* cDNA was PCR amplified from the clone RH09162 obtained from the *Drosophila* Genomics Resource Center, Indiana University, USA, using the following primers with added restriction sites: P1-EcoRI (forward) 5’-CCATCCGAATTCAAAATGCAGCTGATTCTCGT and P2-XhoI (reverse) 5’-TGGCTGCTCGAGATCTATTGGGTTATGAGAGA. The 1160 bp PCR fragment was inserted into pUAST (Brand and Perrimon, 1993), verified by sequencing (GATC Biotech) and sent to BestGene, Chino Hills, CA, USA, for P-element transformation by random insertion in *w*^1118^ background. A 2d-chromosome insertion of *UAS-CG31030* that yielded strong expression of the transgene was used thereafter.

### Reverse transcription-coupled qPCR

Total RNAs were extracted from 20 heads (or 15 thoraces or 15 abdomens) of 8-day-old flies using the QIAzol Lysis reagent (Qiagen). The Maxima First Strand cDNA Synthesis Kit (Thermo Fisher Scientific, ref. K1671) was used with oligo(dT)20 primers to synthesize the cDNAs. Relative quantitative PCR assays were carried out using a LightCycler 480 and the SYBR Green I Master mix (Roche LifeScience), with *Act5C* as internal control for normalization of mRNA levels. All reactions were performed in triplicate. The specificity of amplification products was assessed by melting curve analyses. The following forward and reverse primers, were used: for *CG31030*, 5’-GGCTTCGTTGTAGGCCAACAGA and 5’-CACCAGGTATCCCAAGTTCCAGA; for *Act5C*, 5’-CGTCGACCATGAAGATCAAG and 5’-TTGGAGATCCACATCTGCTG.

### CRISPR/Ca9 gene tagging

The sequence of a V5 tag was inserted in frame after the coding sequence of the *CG31030* gene, using a homology-directed repair CRISPR-Cas9 method (see Figure 1C). The following guide RNA sequence: 5’-TTCACCGTACAGGAGTAAGG-3’ was cloned into the BbsI site of pCFD3: U6:3-gRNA plasmid (Port et al., 2014) (kind gift of Hervé Tricoire, Université de Paris, Paris, France). This plasmid was then injected, at a concentration of 500 ng/μL, with the following single-stranded oligodeoxynucleotide (ssODN) donor repair template: 5’-GTTCGCGCAGCAAACAGTTGACCTTCACCGTACAGGAGTAC<u>GCAGGTAAGCCTATCCCTAACCCTCTCCT CGGTCTAGATTCTACG</u>TAAGGAGGTCATAAGTCTCTGATGAACCAATAGATCTCGGGC-3’ (synthetized by Integrated DNA Technology, Leuven, Belgium), also at a concentration of 500 ng/μL (the sequence underlined corresponds to the in-frame V5 tag), into *nos-cas9* embryos (genotype *y^1^, P(nos-cas9, w+), M(3xP3-RFP.attP)ZH-2A, w\**) (Port et al., 2014). An alanine was added before the V5 tag (dashed underline) to prevent the creation of a potential tyrosine phosphorylation site. Embryo injections were performed by BestGene (Chino Hills, CA, USA). Single F_0_ flies were crossed over the *TM6C*(*Sb*) balancer to establish stable lines. DNA was then extracted from three flies of each of these independent lines, and V5 insertion events were detected by dot blot using a mouse anti-V5 tag monoclonal antibody (Thermo Fisher Scientific, Ref. R960-25). Positive strains were outcrossed in a *w*^1118^ background and their genomic DNA was sequenced to check for proper in-frame V5 integration in *CG31030*. One of these equivalent *CG31030*^V5^ mutant line was selected for further studies.

### Immunohistochemistry

Adult brains of 8-day-old females, or 3^rd^-instar larva CNS, were dissected in *Drosophila* Ringer’s solution or hemolymph-like saline solution (HL3) (70 mM NaCl, 5 mM KCl, 1.5 mM CaCl_2_, 70 mM MgCl_2_, 10 mM NaHCO_3_, 115 mM sucrose, 5 mM trehalose, 5 mM HEPES, with pH adjusted to 7.6), respectively, and fixed in 4% paraformaldehyde (Thermo Fischer Scientific) for one hour. After three 20-min washes in PBS plus 0.5% Triton X-100 (PBT), brains were blocked in PBT + 2% bovine serum albumin for two hours. They were then incubated in primary antibodies diluted in blocking solution for 24 h at 4°C. The primary antibodies used were: mouse monoclonal anti-V5 (Thermo Fisher Scientific, ref. R960-25, 1:200) and rat anti-CadN (DSHB, ref.DN-Ex #8, 1:20). Brains were then washed three times, for 20 min each, in PBT before being incubated in secondary antibodies for 2 hours. Secondary antibodies used were: Alexa Fluor 488 anti-mouse and Alexa Fluor 555 anti-rat (Fisher Scientific, ref.A11029 and ref. A21434 respectively), all diluted at 1:1000. After two 20-min washes in PBT followed by two 20-min washes in PBS, brains were mounted in Prolong Gold Antifade Mountant (Thermo Fisher Scientific, ref. P36930). Imaged were acquired on a Nikon A1R confocal microscope.

For immunostaining of the larval muscles and neuromuscular junctions, the Alexa Fluor 488 Tyramide SuperBoost Kit (Thermo Fisher, ref. B40912) was used to increase the V5 signal that was otherwise faint in this tissue. The working protocol was as recommended by the manufacturer, with anti-V5 diluted to 1:100, then followed by classical immunostaining, as described above, to co-stain for the nerve terminals with an anti-HRP antibody (Jackson ImmunoResearch, ref. 323-005-021, 1:200).

### Longevity assay

About 110 virgin females from each genotype were collected on their hatching day, and placed in clean bottles, with no more than 25 flies per bottle. Flies were transferred in new clean bottles, and survivors were counted every two days for 60 days.

### Co-immunoprecipitation

About 200 heads from 8 day-old *CG31030*^V5^ and *w*^1118^ flies were lysed using glass beads in 500 µL of ice-cold lysis buffer: 50 mM Tris-HCl pH 7.4, 2 mM EDTA, 150 mM NaCl, 0.5% (vol/vol) IGEPAL CA-630 (Sigma-Aldrich, ref. I3021), 10% (vol/vol) glycerol, 1 mM PMSF protease inhibitor (Sigma-Aldrich, ref. P-7626) and 1X cOmplete Mini Protease Inhibitor Cocktail (Roche, ref. 11836153001). Samples were left on ice, with occasional gentle agitation, for 30 min before being centrifuged at 12,000 rpm (13,000 g) at 4°C for 10 min to remove insoluble material. 400 µL of the supernatants were then added to 50 µL of Anti-V5-tag mAb-Magnetic Beads (MBL International, Woburn, MA, USA, ref. M167-11), that had been previously washed as recommended by the manufacturer. Samples were incubated with gentle agitation at 4°C for 4 hours. The supernatants were removed using a magnetic rack, and beads were washed three times with 500 µM of ice-cold lysis buffer before being resuspended in 10 µL of milliQ water.

### Mass spectrometry

The MS/MS analysis was performed at the Proteomics facility of the Institut Jacques Monod (Université de Paris, Paris, France). Briefly, proteins were directly digested on the beads by trypsin (Promega, Madison, WI, USA) overnight at 37°C in a 25-mM NH_4_HCO_3_ buffer (0.2 µg trypsin in 20 µL). The resulting peptides were desalted on a ZipTip μ-C18 Pipette Tips (Pierce Biotechnology, Rockford, IL, USA). Eluates were analyzed using either an Orbitrap Fusion or an Orbitrap Q-Exactive Plus, coupled respectively to a Nano-LC Proxeon 1200 or a Nano-LC Proxeon 1000, both equipped with an easy spray ion source (Thermo Fisher Scientific, Waltham, MA, USA). Raw data were processed on Proteome Discoverer 2.4 with the mascot node (Mascot version 2.5.1), with a maximum of 2 missed cleavages authorized, against the Swissprot/TrEMBL protein database release 2019_12 for *Drosophila melanogaster*. The following post-translational modifications were included as variable: Acetyl (Protein N-term), Oxidation (M), Phosphorylation (STY). Peptide identifications were validated with a 1% FDR (false discovery rate) threshold calculated with the Percolator algorithm. Label-free quantification was done in TOP 3 abundance calculation mode with pairwise ratio-based calculation and t-test (background based) hypothesis test. Only proteins identified in at least one group in two independent experiments were kept in the analysis. Missing values were set to the minimum abundance of the experiment. A more detailed description of the LC-MS/MS procedure is provided in the Supplementary Information.

### VMAT-pHluorin experiments

Third-instar larvae expressing VMAT-pHluorin with or without *CG31030* RNAi in motor neurons using the *OK371-Gal4* driver, were dissected to expose the body wall muscles in Ca^2+^-free HL3 saline solution (70 mM NaCl, 5 mM KCl, 70 mM MgCl_2_, 10 mM NaHCO_3_, 115 mM sucrose, 5 mM trehalose, 5 mM HEPES, pH 7.6). Two other solutions were used: an acidic Ca^2+^-free HL3 saline (pH 5.5) and a neutral HL3-NH_4_Cl saline (pH 7.6), in which 50 mM NaCl was replaced by 50 mM NH_4_Cl. After dissection, larvae fillets were allowed to settle in Ca^2+^-free HL3 saline for 10 min before being scanned a first time directly in a drop of the solution using a Nikon A1R confocal microscope. Ca^2+^-free HL3 saline was then replaced either by the acidic Ca^2+^-free HL3 saline for low pH-induced quenching experiments, or by the HL3-NH_4_Cl saline for pH gradient collapse experiments. In both cases, larval fillets were rinsed three times with the modified solution, and incubated for 3 min, before being scanned a second time, in a drop of modified solution. Quantification was done on Z-projections (set to maximal intensity) of confocal stacks. Using the Fiji software, a fixed threshold was applied to all images to get rid of the background and select only synaptic areas. Percentage of area over the threshold was used as a measure of the signal intensity and ratios of the values obtained in modified solutions over standard HL3, i.e. the second scan over the first scan, were calculated for quantifications.

### Larval locomotion assays

Larval locomotion was assessed on an in-house made version of the FIMtable system (Risse et al., 2014). Third-instar larvae were collected and briefly rinsed in water to remove traces of food, before being gently placed on the recording table precoated with a thin layer of 1.2% agar gel. Only four larvae of the same genotype were recorded simultaneously, to avoid collisions between animals, for a period of 2 min with a Basler ace acA2040-25gm camera at 12.5 frames/s. Larvae that burrowed themselves into the agar plate or escaped the arena before the end of recording were excluded from the results. Tracking was done using the FIMtrack software, as described in (Risse et al., 2014). The number of peristaltic waves was computed from the variations in larval area. More precisely, the curves of area variation were first smoothed with a Savitzky-Golay filter to get rid of unwanted noise, then the number of waves was defined as the number of peaks on the curve (which was automatically computed by a custom-made Python script). Stride size was then calculated as the distance travelled by the larva divided by the number of peristaltic waves, while stride duration was defined as the recording time (i.e. 120 seconds) divided by the number of peristaltic waves.

### Electrophysiological recording and quantal analysis

Third-instar larvae expressing *CG31030* RNAi in motor neurons using the *OK371-Gal4* driver, and appropriate controls, were dissected to expose the body wall muscles, and the brain removed, in Ca^2+^-free HL3 saline solution (70 mM NaCl, 5 mM KCl, 70 mM MgCl_2_, 10 mM NaHCO_3_, 115 mM sucrose, 5 mM trehalose, 5 mM HEPES, pH 7.6) (Stewart et al., 1994; Cattaert and Birman, 2001). Spontaneous miniature EPSPs (mEPSPs) were recorded in the presence of tetrodotoxin (TTX) 10^−6^M, so that no spike could occur, from the ventral longitudinal abdominal muscle 6 in segment A3. Quantal analysis was performed following the theoretical background described in (Kuno, 1971) and (Castellucci and Kandel, 1974). Distribution histograms of mEPSP size were built from each muscle fiber recording with a 0.01 mV bin size. These histograms provided an estimate of the mean size of a unitary EPSP, since the peaks represent integer multiples of the unitary size. A theoretical distribution was then computed by convolving a binomial distribution, accounting for quantal content (number of quanta released), and a Gaussian distribution, allowing for variations in size of individual quanta. A detailed description of the mathematical treatment is provided in the Supplementary Information.

### Statistical Analysis

Statistical analysis was performed with the GraphPad Prism 6 software. The paired Student’s t-test was used for comparison of two genotypes, while either paired or unpaired ANOVA, with Dunnett’s post-test for multiple comparisons, were used for three genotypes.

## Acknowledgements

We thank Dr Philippe Marin for helpful advice about the quantitative analysis of mass spectrometry experiments, Drs Hervé Tricoire and Elodie Martin for helpful suggestion on the CRISPR/Cas9 mutagenesis, Pr David Krantz for providing the *UAS-VMAT-pHluorin* strain, Dr Hervé Tricoire for the *pCFD3: U6:3-gRNA* plasmid, and Dr. Céline Petitgas and Manon Dobrigna for participating in preliminary experiments during their student internships. This work was supported by a grant from the Fondation de France (Subvention n° 0086407) to SB and recurrent fundings to his lab from the ESPCI Paris and CNRS. AD, ARI and JS were recipient of PhD fellowships from PSL University and Labex MemoLife, the Doctoral School 3C and the China Scholarship Council, respectively.

## Supplementary information

### Supplementary figures

**Supplementary Figure 1.**
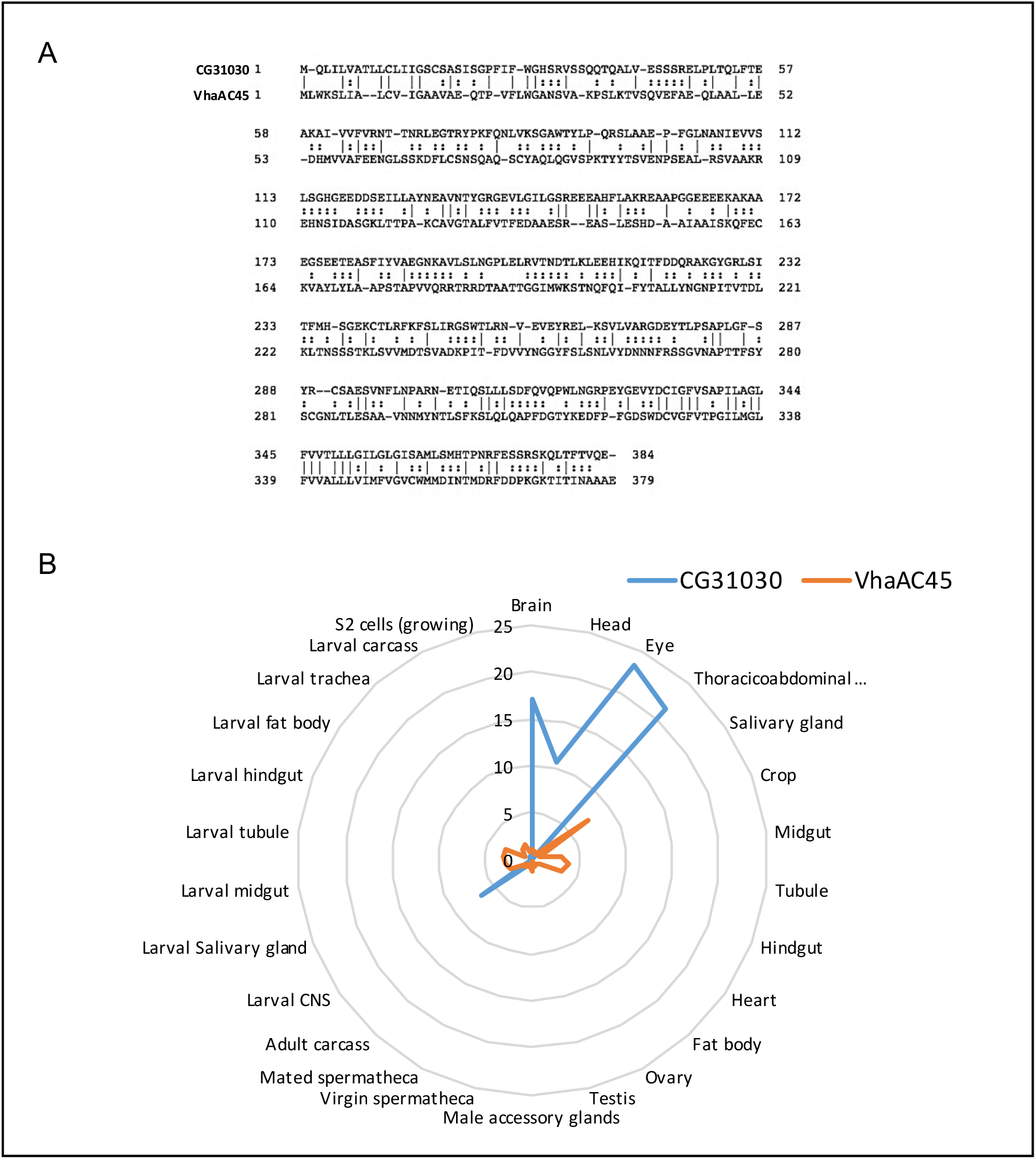
*CG31030* is a paralog of *VhaAC45* expressed in the nervous system. (**A**) Amino-acid sequence alignment shows that CG31030 and VhaAC45 share 69.9% similarity. (B) Diagram representing expression levels of *CG31030* and *VhaAC45* in different tissues relative to the whole fly, according to FlyAtlas data (Chintapalli et al., 2007). CG31030 appears to be markedly enriched in the nervous system of larva and adult fly, whereas in contrast VhaAC45 seems to be uniformly expressed in all tissues in these two stages.

**Supplementary Figure 2.**
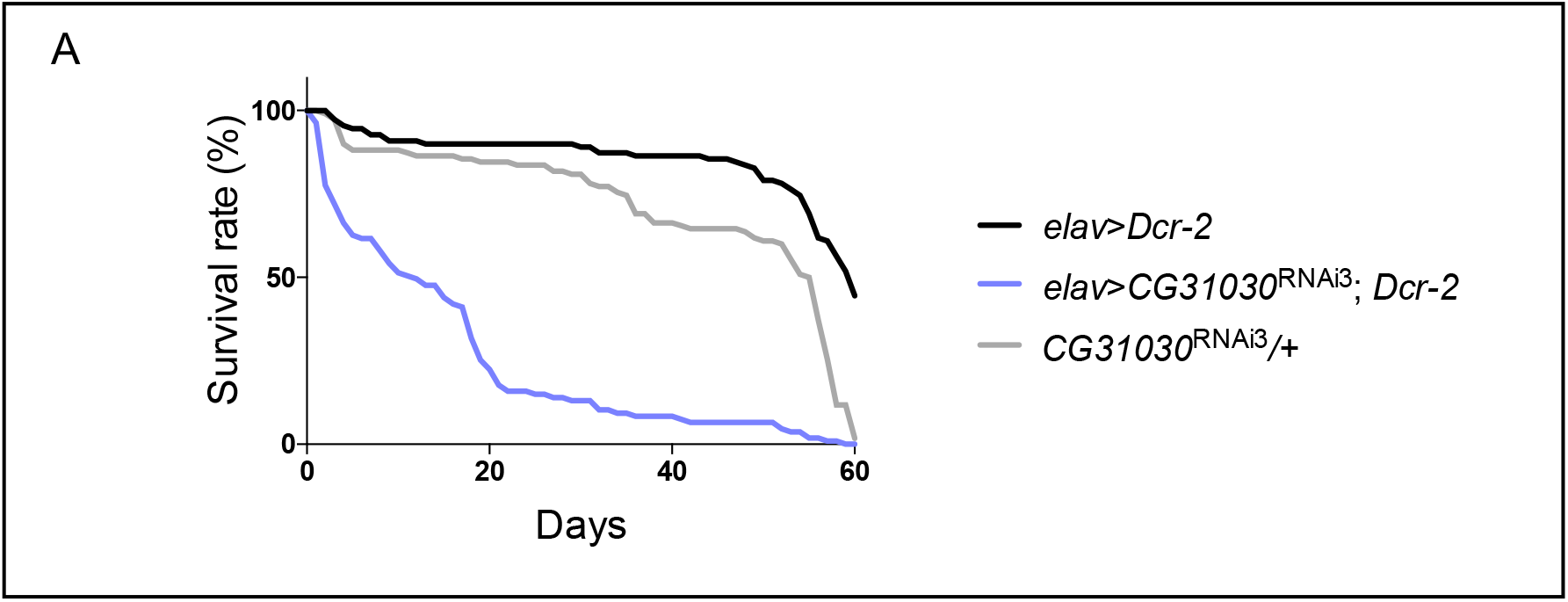
Knockdown of *CG31030* expression in neurons decreases adult longevity. Pan-neuronal expression of *CG31030*^RNAi3^ and *Dcr-2* with the *elav-Gal4* driver led to a marked shortening of the lifespan of adult flies. Results of one experiment, carried out with 105-110 females per genotype.

**Supplementary Figure 3.**
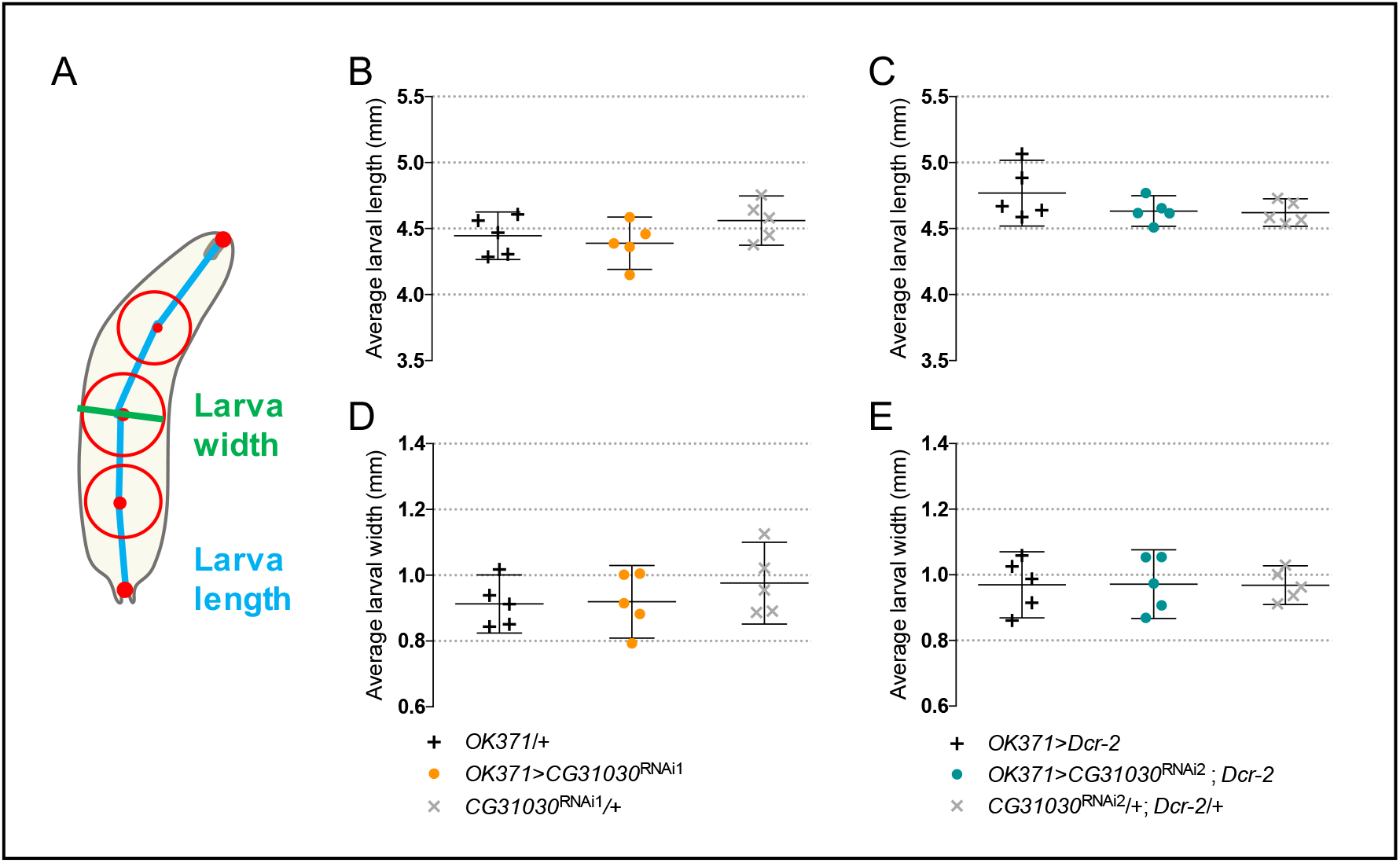
CG31030 downregulation in motoneurons did not significantly alter larval size. (**A**) Larval length is defined as the spine length from head to tail, while larval width is the diameter of the mid-spine circle. (**B-E**) Average larval length (**B**, **C**) and width (**D**, **E**) were not significantly different between animals expressing *CG31030* RNAi in motoneurons and controls.

### Supplementary tables

**Supplementary Table 1.** Pan-neuronal expression of *CG31030* rescues the embryonic lethality of CG31030-deficient flies up to the adult stage.

| Genotype of the progeny flies<sup>1</sup> | Expected percentage<sub>2</sub> | Scored percentage<sub>3</sub> |
| --- | --- | --- |
| Heterozygous deficiency without *CG31030* expression | 40 | 39.7 |
| Heterozygous deficiency with pan-neuronal *CG31030* expression | 40 | 53.8 |
| Homozygous deficiency without *CG31030* expression | 0 | 0 |
| Homozygous deficiency with pan-neuronal *CG31030* expression | 20 | 6.4 |

<sup>1</sup>Rescue of embryonic lethality was assayed by crossing *elav-Gal4; CG31030<sup>MI107</sup>/TM6B(Tb)* females with *w; UAS-CG31030/CyO; Df(3R)Exel6214/TM6B(Tb)* males, and scoring the relative number of *CG31030*-expressing adult flies homozygous for *CG31030* deficiency (i.e. non-*Tb* and non-*Cy elav-Gal4/+; UAS-CG31030/+; CG31030<sup>MI107</sup>/Df(3R)Exel6214*) in the progeny (highlighted line).
<sup>2</sup>Expected percentage of adult progeny flies of each genotype in case of full rescue.
<sup>3</sup>Actual percentage obtained in the experiments. 24 rescued adults were recovered out of a total of 373 progeny flies.

**Supplementary Table 2.** List of the genes encoding proteins that co-immunoprecipitated with CG31030 in *Drosophila* head extracts.

| Gene Symbol<sup>1</sup> | FlyBase Gene ID | Log<sub>2</sub>(R<sub>1</sub>) | Log<sub>2</sub>(R<sub>2</sub>) | Log<sub>2</sub>(R<sub>3</sub>) |
| --- | --- | --- | --- | --- |
| *ATP6AP2* | FBgn0037671 | 6.154281057 | 9.481786693 | 5.17436482 |
| *Vha100-1* | FBgn0028671 | 1.499248537 | 8.012982519 | 5.43599503 |
| *VhaAC39-1* | FBgn0028665 | 2.104797507 | 8.293264658 | 4.86741949 |
| *CG31030* | FBgn0051030 | 3.669278876 | 7.960835791 | 5.19465574 |
| *Twdlβ* | FBgn0033658 | 1.642952792 | 3.068640973 | 2.35207192 |
| *Ccp84Ag* | FBgn0004777 | 1.543972087 | 1.326485589 | 1.86392045 |
| *CG13627* | FBgn0039217 | 5.738384187 | 4.008318188 | 3.18433108 |
| *mfas* | FBgn0260745 | 1.596554441 | 3.368384477 | 3.03376685 |
| *CG16820* | FBgn0032495 | 4.77051909 | 9.784086057 | 3.60280541 |
| *Cpr64Ab* | FBgn0035511 | 1.08994458 | 2.758609792 | 2.31103817 |
| *CG14752* | FBgn0033307 | 2.21698759 | 3.69458382 | 3.10154816 |
| *CG15615* | FBgn0034159 | 2.77101281 | 5.12362053 | 3.39808341 |

<sup>1</sup>12 proteins were identified with at least twice higher abundance in *CG31030*<sup>V5</sup> compared to the *w*<sup>1118</sup> control in three independent co-immunoprecipitations with anti-V5 antibodies, followed by mass-spectrometry analysis experiments. Among these proteins, three are known to be constitutive or accessory subunits of the V-ATPase complex. They are indicated in the first ochre-highlighted lines of the Table and circled in blue in Fig. 2. The grey line indicates the co-immunoprecipitation target CG31030.

**Supplementary Table 3.**
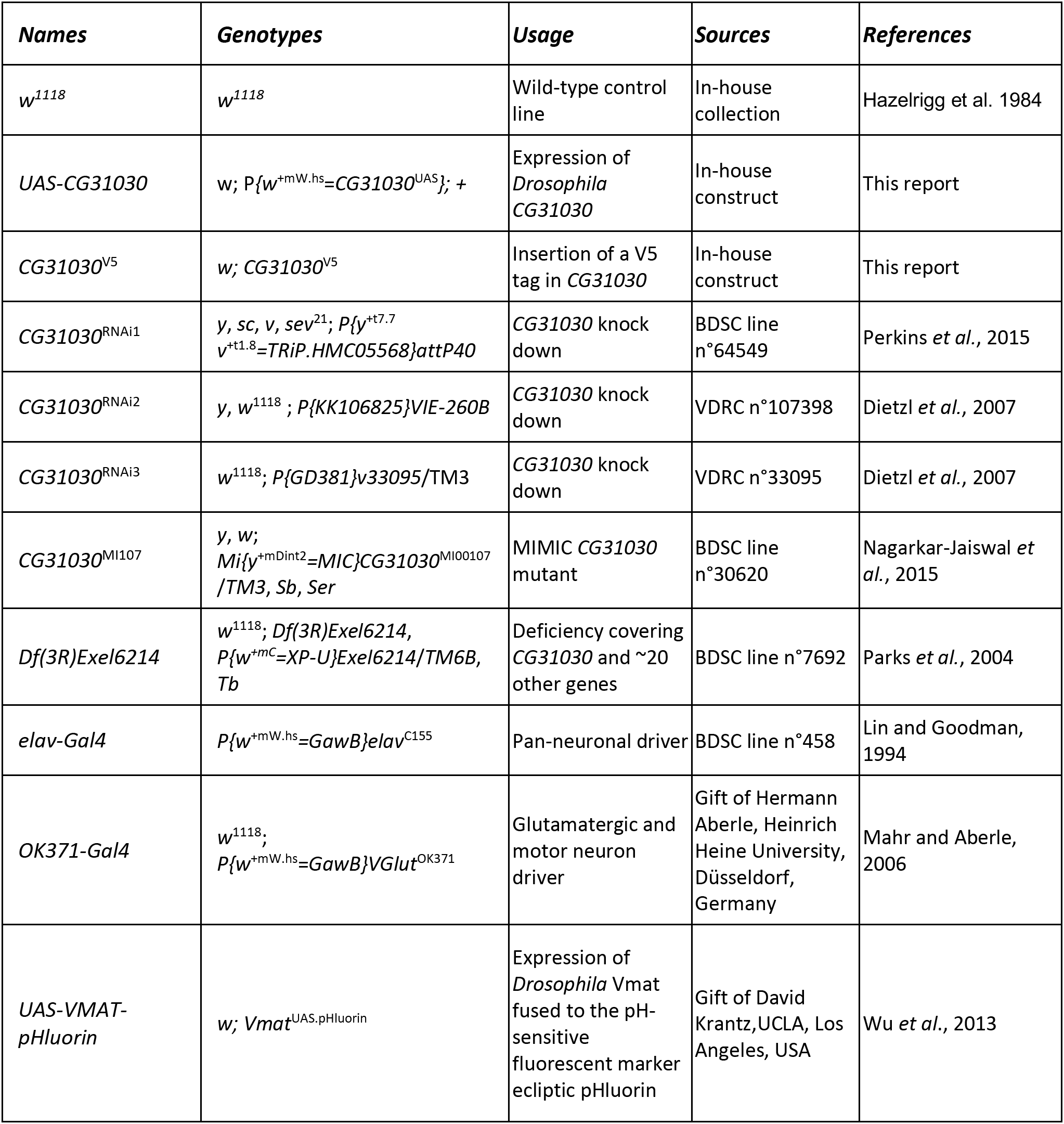
*Drosophila* strains used in this study.

### Supplementary methods

#### Detailed LC-MS/MS procedure

Proteins on beads were digested overnight at 37°C with trypsin (Promega, Madison, WI, USA) in 25 mM NH_4_HCO_3_ buffer (0.2 µg trypsin in 20 µL). The resulting peptides were desalted using ZipTip μ-C18 Pipette Tips (Pierce Biotechnology, Rockford, IL, USA). Samples were analyzed using either an Orbitrap Fusion or an Orbitrap Q-Exactive Plus, coupled respectively to a Nano-LC Proxeon 1200 or a Nano-LC Proxeon 1000, both equipped with an easy spray ion source (Thermo Fisher Scientific, Waltham, MA, USA). On the Orbitrap Fusion instrument, peptides were loaded with an online preconcentration method and separated by chromatography using a Pepmap-RSLC C18 column (0.75 x 750 mm, 2 μm, 100 Å) from Thermo Fisher Scientific, equilibrated at 50°C and operated at a flow rate of 300 nl/min. Peptides were eluted by a gradient of solvent A (H_2_O, 0.1 % FA) and solvent B (ACN/H_2_O 80/20, 0.1% FA). The column was first equilibrated for 5 min with 95 % of A, then B was raised to 28 % in 105 min and to 40% in 15 min. Finally, the column was washed with 95% B during 20 min and re-equilibrated at 95% A for 10 min. Peptides masses were analyzed in the Orbitrap cell in full ion scan mode, at a resolution of 120,000, a mass range of *m/z* 350-1550 and an AGC target of 4.10^5^. MS/MS were performed in the top speed 3s mode. Peptides were selected for fragmentation by Higher-energy C-trap Dissociation (HCD) with a Normalized Collisional Energy of 27% and a dynamic exclusion of 60 seconds. Fragment masses were measured in an Ion trap in the rapid mode, with and an AGC target of 1.10^4^. Monocharged peptides and unassigned charge states were excluded from the MS/MS acquisition. The maximum ion accumulation times were set to 100 ms for MS and 35 ms for MS/MS acquisitions respectively.

On the Q-Exactive Plus instrument, peptides were loaded with an online preconcentration method and separated by chromatography using a Pepmap-RSLC C18 column (0.75 x 500 mm, 2 μm, 100 Å) from Thermo Scientific, equilibrated at 50°C and operated at a flow rate of 300 nl/min. Peptides were eluted by a gradient of solvent A (H_2_O, 0.1 % FA) and solvent B (100 % ACN, 0.1% FA), the column was first equilibrated 5 min with 95 % of A, then B was raised to 35 % in 93 min and finally, the column was washed with 80% B during 10 min and re-equilibrated at 95% A during 10 min. Peptides masses were analyzed in the Orbitrap cell in full ion scan mode at a resolution of 70,000 with a mass range of *m/z* 375-1500 and an AGC target of 3.10^6^. MS/MS were performed in a Top 20 DDA mode. Peptides were selected for fragmentation by Higher-energy C-trap Dissociation (HCD) with a Normalized Collisional Energy of 27%, and a dynamic exclusion of 30 seconds. Fragment masses were measured in the Orbitrap cell at a resolution of 17,500, with an AGC target of 2.10^5^. Monocharged peptides and unassigned charge states were excluded from the MS/MS acquisition. The maximum ion accumulation times were set to 50 ms for MS and 45 ms for MS/MS acquisitions respectively.

Raw data were processed on Proteome Discoverer 2.4 with the mascot node (Mascot version 2.5.1), with the Swissprot/TrEMBL protein database release 2019_12 for *Drosophila melanogaster*. A maximum of 2 missed cleavages was authorized. Precursor and fragment mass tolerances were set to respectively 7 ppm and 0.5 Da (Orbitrap Fusion) and to 6 ppm and 0.02 Da (Orbitrap Q-exactive Plus). The following post-translational modifications were included as variable: Acetyl (Protein N-term), Oxidation (M), Phosphorylation (STY). Spectra were filtered using a 1% FDR (false discovery rate) with the percolator node. Label-free quantification was done in TOP 3 abundance calculation mode with pairwise ratio based calculation and t-test (background based) hypothesis test.

#### Quantal analysis

The probability for an EPSP to contain *i* quanta followed a binomial distribution:

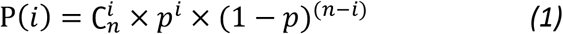

in which *p* is the average probability of release, and *n* is the total number of releasable quanta. In the frequency distribution of EPSP amplitude, successive peaks represent increasing numbers of quanta.

In order to predict how these events are distributed between different bins in a histogram, it is necessary to allow for variations in quantal size. To do this, the largest peaks of the histogram were fitted to a Gaussian curve scaled in width to have a variance proportional to quantal size, and scaled in height so that its area corresponded to the predicted number of events (Del Castillo and Katz, 1954). From the mean (*μ*) and standard deviation (*σ*) of the Gaussian curve, the content (*f*) of each bin (*y*) is given by:

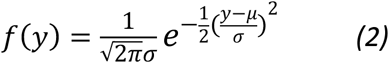

The standard deviation (*σ*) of each peak depends on the number of quanta it contains. It was calculated as follows:

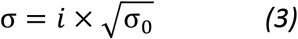

with *i* the number of quanta in the peak, and σ_0_ the standard deviation of a single quanta. The amplitude of Gaussian distribution for each peak is scaled to the probability *P*(*i*) for that peak (see *(1)*).

The complete theoretical distribution, allowing for variance and peak overlap was then obtained by pooling the Gaussians for all peaks. In this way, the theoretical distribution could be superimposed on the histogram for direct comparison with the data.

